# Resilience in soil bacterial communities of the boreal forest from one to five years after wildfire across a severity gradient

**DOI:** 10.1101/2022.04.21.487669

**Authors:** Thea Whitman, Jamie Woolet, Miranda Sikora, Dana B. Johnson, Ellen Whitman

## Abstract

Wildfires can represent a major disturbance to ecosystems, including soil microbial communities belowground. Furthermore, fire regimes are changing in many parts of the world, altering and often increasing fire severity, frequency, and size. The boreal forest and taiga plains ecoregions of northern Canada are characterized by naturally-occurring stand-replacing wildfires on a 40-350 year basis. We previously studied the effects of wildfire on soil microbial communities one year post-fire across 40 sites, spanning a range of burn severity. Here, we return to the same sites five years post-fire to test a series of hypotheses about the effects of fire on bacterial community composition. We ask the following questions: (1a) Do the fundamental factors structuring bacterial community composition remain the same five years post-fire? (1b) Do the effects of fire on bacterial community composition decrease between one and five years post-fire? (1c) Do shifts in bacterial community composition between one and five years post-fire suggest resilience? (2a) Does the importance of fast growth diminish between one and five years post-fire? (2b) Do short-term post-fire responders continue to dominate the community five years post-fire? We find the following: (1a) Five years post-fire, vegetation community, moisture regime, pH, total carbon, texture, and burned/unburned all remained significant predictors of bacterial community composition with similar predictive value (R^2^). (1b and 1c) Bacterial communities became more similar to unburned sites five years post-fire, across the range of severity, suggesting resilience, while general structure of co-occurrence networks remained similar one and five years post-fire. (2a) Fast growth potential, as estimated using predicted 16S rRNA copy numbers, was no longer significantly correlated with burn severity five years post-fire, indicating the importance of this trait for structuring bacterial community composition may be limited to relatively short timescales. (2b) Many taxa that were enriched in burned sites one year post-fire remained enriched five years post-fire, although the degree to which they were enriched generally decreased. Specific taxa of interest from the genera *Massilia, Blastococcus*, and *Arthrobacter* all remained significantly enriched, suggesting that they may have traits that allow them to continue to flourish in the post-fire environment, such as tolerance to increased pH or ability to degrade pyrogenic organic matter. This hypothesis-based work expands our understanding of the post-fire recovery of soil bacterial communities and raises new hypotheses to test in future studies.

## 1. Introduction

Wildfires burn an estimated 300-400 million ha of land globally each year (Lierop et al., 2015), releasing CO_2_ emissions equivalent to roughly half those of fossil fuels (Bowman et al., 2009). Belowground, the biogeochemical aftermath of wildfire continues long after the burn: in boreal and temperate forest fires, it takes on the order of one hundred years for soil C and N stocks to recover (Nave et al., 2011; Mack et al., 2021), and large pulses of mineral N are often released in the months following fire (Wan et al., 2001). These dynamics are mediated by surviving and recolonizing soil microbes (Smithwick et al., 2005), which also interact with plant recolonization post-fire (Knelman et al., 2015). Understanding how wildfires affect soil microbial processes and ecosystem responses to disturbance will require an understanding of not only which microbes respond to fires, but, more importantly, why they do so. For fungi, response to fire has been studied scientifically for over a century (Boudier, 1877; Seaver, 1909), and human knowledge of certain “pyrophilous” fungi, such as certain morels, is likely ancient (Anderson and Lake, 2013). For plants, fire ecology frameworks are richly developed and well-established (Cooper, 1961). However, our understanding of how bacteria respond to fires is only beginning to emerge.

Fires can directly reduce total soil microbial biomass and often result in changes to community composition and decreased microbial diversity (Dooley and Treseder, 2011; Pressler et al., 2018; Certini et al., 2021). It can take years to observe resilience within soil microbial communities after a disturbance (considering resilience as a return to similar pre-burn community composition) (Allison and Martiny, 2008), and may not be achieved for decades after wildfires (Dooley and Treseder, 2011; Holden and Treseder, 2013; Köster et al., 2014). As Shade et al. (2012) note, time and spatial scale affect whether a given disturbance might be considered a “pulse” or a “press” disturbance (*sensu* (Bender et al., 1984)). Wildfires, at the spatial and temporal scales that are relevant to soil microbes, could perhaps be categorized as hybrid pulse / press disturbances (Bender et al., 1984). The immediate effects of wildfire – death via combustion and high temperatures – are a classical pulse disturbance, but the subsequent shifts in soil conditions, such as loss of the organic (O) horizon, transformation of organic matter, or increase in pH, persist, eliciting a press disturbance. Fire severity would moderate the effect of each of these types of disturbances on soil microbes, with both expected to intensify with increasing severity. In the case of high severity forest fires, where the O horizons can be completely combusted, a long time before recovery is unsurprising – the habitat of O horizon-dwelling microbes has literally been destroyed, and so it is reasonable to expect that a full return to the community’s previous state would require the re-accumulation of this organic horizon, which can take decades (Lecomte et al., 2006; Andrieux et al., 2018). In mineral soil horizons, the habitat itself (i.e., mineral particles) largely remains, but can be severely altered by high temperatures and elemental transformations (e.g., organic matter combustion, pyrogenic organic matter production, volatilization of some elements, and deposition of others) (Certini, 2005). Given the strong role that fire-affected soil properties such as pH and organic matter content play in structuring microbial community composition (Bahram et al., 2018; Delgado-Baquerizo et al., 2018), it is also not surprising that a return to pre-fire soil microbial community compositions might occur on timescales at least as long as those required for key community-structuring soil properties to return to their pre-fire state. We observed similarly strong linkages when examining wildfire effects on soil microbial community composition along a gradient of fire severity (Whitman et al., 2019).

Fire severity (sometimes called burn severity) can be quantified or estimated using a range of different metrics, depending on the ecosystem, study questions, and management goals, but we will use the term fire severity here *sensu* Keeley (2009) – *i*.*e*., degree of loss of organic matter above- and belowground. Fire severity relates to both fire intensity (energy released during the fire) and the degree of ecological impact from the fire. Fire severity varies across ecosystems – e.g., frequent, lower-severity fires characteristic of grasslands of the Midwest United States vs. less frequent higher-severity fires characteristic of boreal forests of northern Canada. Fire severity is also heterogeneous across the landscape within a single fire (Holden et al., 2016). A number of recent studies have quantified the effects of wildfires on soil bacterial communities across a range of fire severity (Weber et al., 2014; Holden et al., 2016; Brown et al., 2019; Lucas-Borja et al., 2019; Whitman et al., 2019; Adkins et al., 2020; Miera et al., 2020), often observing greater changes in microbial community composition with increasing severity. However, most of these studies consider relatively short timescales (3 years or less) and/or single timepoints, which does not allow us to follow the recovery of communities over longer periods of time. While chronosequence-based studies have indicated the range of timescales required for community composition and soil microbial functions to return to pre-burned states (Sun et al., 2015, 2016; Pérez-Valera et al., 2018, 2020; Zhou et al., 2020), there are notable assumptions required for and limitations to space-for-time study designs. More importantly, quantitatively assessing burn severity at time of fire in a chronosequence study can be difficult, since most sites, by definition, are somewhere along a trajectory of recovery, and post-hoc assessments of burn severity are difficult. Thus, as noted by Pérez-Valera et al. (2020), the question of the degree to which fire severity affects bacterial community resilience remains to be thoroughly investigated. A need for long-term post-fire soil biological community monitoring is also highlighted by Certini et al. (2021), as a key way to better understand the specific ecology of individual fire-responsive taxa.

Although the field is currently in a nascent stage compared to plant ecology and perhaps even fungal ecology, we are beginning to gain a sense for the fire ecology of specific soil bacteria beyond observed changes at the whole-community level. Support for the importance of certain “pyrophilous” organisms and their associated traits is beginning to emerge. For example, some taxa from the genus *Arthrobacter* increase in relative abundance by orders of magnitude after wildfires across different ecosystems (Weber et al., 2014; Fernández-González et al., 2017; Whitman et al., 2019; Miera et al., 2020). We can propose characteristics that may help promote these positive fire responders, such as spore-forming capabilities possibly supporting fire survival (Mongodin et al., 2006), the ability to grow quickly post-fire (Nemergut et al., 2016; Whitman et al., 2019), or the capacity to mineralize fire-altered organic matter (Westerberg et al., 2000). These and other pyrophilous traits may map on to equivalent plant traits in some cases – e.g., bacteria that survive fires via spores or other means represent an analogue to plant seed banks, where rapid regeneration or resprouting post-fire can strongly influence subsequent communities (Johnstone et al., 2016) – and not in other cases – e.g., the ability to use pyrogenic organic matter (PyOM) as a C source does not have a clear analogue in plants. It is important to note these inferences often remain somewhat speculative, however, since conclusively demonstrating functions or traits from amplicon sequencing data – currently the most common method of community characterization – is problematic. Additionally, the relative importance of each of these traits would be expected to vary over time as post-fire succession proceeds. As with whole-community metrics, most of what we have inferred about fire ecology for individual bacterial taxa is again based on short term and/or single timepoint studies. In the current study, we seek to determine how wildfires affect soil bacterial communities and individual taxa over time, with an emphasis on how fire severity affects community resilience and the relative importance of putative pyrophilous traits.

We investigate these questions in an ongoing field study of the widespread wildfires in the boreal forest of northern Canada in 2014, including 40 sites distributed across a range of burn severities and vegetation communities, where we have sampled organic and top mineral soil horizons one year and five years post-fire. We had a series of specific questions and hypotheses that we set out to test, informed by our findings from one year after the burns (reported in (Whitman et al., 2019)):

### Theme 1. Which factors control bacterial community composition during post-fire recovery?

1a. Do the fundamental factors structuring bacterial community composition shortly after a fire remain the same five years post-fire? Specifically, how will significant predictors of community composition and the co-occurrence network change between one and five years post-fire? First, we predicted that the same significant (non-fire) predictors of bacterial community composition (vegetation community, moisture regime, pH, total C, and texture) would remain the same, five years post-fire, without substantial changes in their predictive value (as measured by R^2^). Second, we predicted that the general structure of the co-occurrence network (as measured by which taxa positively or negatively co-occur) would be preserved, because the same factors that make taxa occur at the same sites or different sites would remain constant. However, because we expected communities across the dataset to exhibit trends toward resilience (1c), as burned sites begin to return to their pre-burn states, we also anticipated that we might detect fewer significant nodes (taxa) and connections (positive or negative co-occurrences) within the network. This prediction is based on the rationale that significant co-occurrences will be harder to detect if community composition is more similar across the full dataset.

1b. Do the effects of fire on bacterial community composition decrease between one and five years post-fire? Specifically, will whether sites were burned or not remain a significant predictor of community composition after controlling for the non-fire predictors above by including them in the statistical model? We predicted that burned/unburned would remain a significant predictor five years post-fire, without substantial changes in its predictive value (as measured by R^2^). Although R^2^ for burned/unburned as a predictor could decrease as communities begin to recover from burning, we also recognized that chronosequence studies have observed that soil bacterial community composition can take decades to return to their pre-burned states (Sun et al., 2015; Pérez-Valera et al., 2018, 2020; Zhou et al., 2020), which is relevant given the long historical stand-replacing fire return interval in this region (40-350 years (Boulanger et al., 2012)). Additionally, we recognized that certain effects of the fire on soil bacterial communities – particularly those mediated by changes to vegetation as it follows successional trajectories (Hart et al., 2005) – might accumulate, rather than diminish, over time, adding to the rationale for our prediction.

1c. Do shifts in bacterial community composition between one and five years post-fire suggest resilience? Specifically, will sites that were more severely burned remain more dissimilar from unburned sites, five years post-fire? While we predicted that the overarching predictive factors of community composition would remain similar (1a and 1b), we predicted that the relationship between burn severity and community dissimilarity from unburned sites would remain significant, but become weaker with time – *i*.*e*., burned sites would all become more similar to unburned sites, exhibiting resilience, with larger shifts toward recovery (shifts toward unburned community composition) in the more severely burned sites. Our rationale was that high severity burns are part of the historical fire regime in this region, so communities subjected to high severity burns would also exhibit resilience over some timescale. Since communities from higher severity burns were more dissimilar from unburned communities one year after the fire, some processes such as recolonization from nearby unburned communities or less-affected subsoils might have greater effects on community composition.

### Theme 2. How does the importance of different fire-responsive traits change during post-fire recovery?

2a. Does the importance of fast growth diminish between one and five years post-fire? Specifically, will weighted mean predicted rRNA gene copy number (a proxy for potential for fast growth (Klappenbach et al., 2000; Roller et al., 2016)) remain significantly and positively correlated with burn severity, five years post-fire? We predicted that potential for fast growth would remain an important factor in structuring the community, although the relationship between rRNA gene copy number and burn severity would be weaker. This is based on the assumption that community-structuring processes other than fast growth would have emerged after five years and would be consistent with the results from Nemergut et al. (2016), who observed a significant reduction in mean rRNA gene copy number between 3 and 29 months post-fire.

2b. Do short-term post-fire responders continue to dominate the community five years post-fire? Specifically, will the same taxa remain enriched in burned sites, and to what degree? We predicted that there would be a similar total number of fire-enriched taxa, but that these taxa would include some novel responders while excluding some of the taxa that were fire-enriched one year post-fire, and that the degree of enrichment would be lower overall, as the community as a whole begins to return to its unburned state. More specifically, we predicted that *Blastococcus*, a strong fire responder identified in numerous ecosystems (Fernández-González et al., 2017; Whitman et al., 2019) and enriched at sites from this same region that had undergone short-interval reburns (Woolet et al., *in review*), might be a canonical short-term “pyrophile”, and thus might be less enriched in burned sites five years post-fire. Conversely, we predicted that previously identified burn responders *Arthrobacter* and *Massilia* OTUs (Whitman et al., 2019), which have putative abilities to decompose polyaromatic hydrocarbons (Westerberg et al., 2000; Liu et al., 2014) such as those that might be created during fires and persist over super-decadal timescales (Czimczik and Masiello, 2007), would remain similarly enriched in burned sites five years post-fire.

## 2. Methods

### 2.1 Study region and site selection

Our study region is in the southern half of the boreal and taiga plains ecoregions of northwestern Canada (northern Alberta and the southern Northwest Territories). The study region has long, cold winters and short, hot summers, with mean annual temperatures between −4.3 °C and −1.8 °C and annual precipitation ranging from 300 to 360 mm (ESWG, 1995; Wang et al., 2012). The region’s fire regime is characterized by infrequent stand-replacing fires every 40-350 years on average (Boulanger et al., 2012) and, due to its small and dispersed population, fires are often managed with little suppression and control, when appropriate. 2014 was an exceptional year for wildfires in the region (Whitman et al., 2018b), and offered an opportunity to study wildfires across a range of vegetation communities and burn severities. We selected 40 sites in the Northwest Territories and northern Alberta (Wood Buffalo National Park), Canada, and sampled them one year post-fire, in 2015, and again five years post-fire, in 2019 (Figure 1). The fires and the drivers of burn severity are described in detail in Whitman et al. (2018b), their effects on understory vegetation are described in detail in Whitman et al. (2018a), and their one-year effects on soil microbial communities are described in detail in Whitman et al. (2019). This study builds on the Whitman et al. (2019) dataset to compare bacterial and archaeal community response one year vs. five years post-fire.

**Figure 1.**
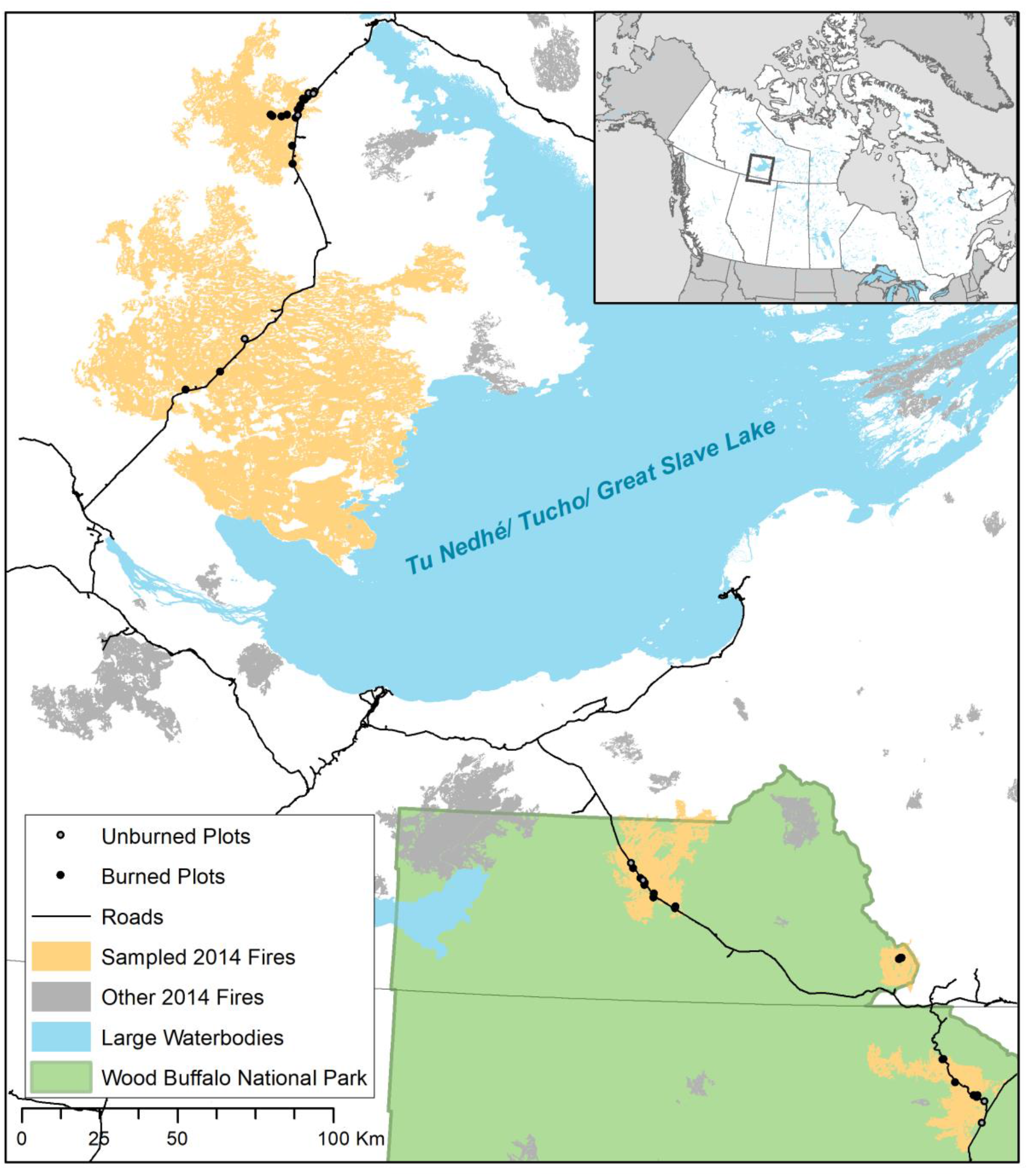
Study region of northern Alberta and the Northwest Territories, Canada, including Wood Buffalo National Park (WBNP – green shading). Closed circles indicate burned plots, open circles indicate unburned plots, yellow shapes indicate sampled fires, and grey shapes indicate other 2014 fires in the region. Inset indicates relative location within Canada.

The six wildfires in this study were very large (14,000 to 700,000 ha). The soils in these regions are mostly classified as Typic Mesisols (21 sites), Orthic Gleysols (6 sites), or Orthic Gray Luvisols (10 sites) (Soil Landscapes of Canada map v.3.2). They span a wide range of soil properties, with pH values ranging from 3.2 (treed wetlands) to 8.1 (uplands with calcareous soil), total C ranging from 0.5% (mineral horizon) to 52% (organic horizon), and a wide range of soil textures. Vegetation communities were classified as jack pine-dominated (*Pinus banskiana* Lamb.) uplands, black spruce-dominated (*Picea mariana* (Mill.)) uplands, a mix of upland coniferous and broadleaf trees, or treed wetlands (Beckingham and Archibald, 1996).

### 2.2 Site assessment methodologies

Sites were selected and characterized as described in detail by Whitman et al. (2018a; 2018b). Briefly, field sites were selected to represent the local range of burn severity and vegetation communities, resulting in a total of 31 burned field sites across three vegetation communities (pine, spruce, and mixedwood). We selected an additional 9 control sites (not burned within the last 38 years before sampling, mean time since fire 95 years; “unburned”), chosen to reflect the range of vegetation communities sampled in the burned plots, for a total of 40 sites. These represent 40 of the 62 original sites (we retained treed wetland sites but did not include open wetlands in this re-analysis, and some sites were not accessible five years post-fire due to a lack of helicopter access, active fires, or closed roads). At each site, one year post-fire, we established a 30 × 30 m square plot with 10 × 10 m subplots at the four corners. We assessed burn severity in the four subplots using burn severity index (BSI (Loboda et al., 2013); described in detail in Whitman et al. (2018a)). We returned to sites five years post-fire to re-assess vegetation composition (Dawe et al. *in review*) and sample soils.

At each plot, we took soil cores (5.5 cm diameter, 13.5 cm depth) at three locations. The sampling scheme was modified between sampling years: at one year post-fire, samples were taken at plot centre, 7 m SW of centre, and 7 m NE of centre, and at five years post-fire, samples were taken at plot centre, 17.5 m N of centre, and 17.5 m S of centre (these changes are not expected to affect our central findings, as indicated by comparing unburned sites between years). Soil cores were gently extruded and separated into organic (O) horizons (where present) and mineral (M) horizons (where present in the top 13.5 cm of soil profile). Where mineral horizons were present, one year post-fire, they were sampled to whatever depth represented the bottom of the 13.5 cm core. Five years post-fire, we modified the protocol so a consistent depth (5 cm) was sampled for mineral soils. (Although the two years’ sampling approaches would have rarely or never included different genetic soil horizons, this would undoubtedly affect certain soil properties that vary with depth. However, we do not expect that this methodological change is driving any of our key findings, as indicated by broad similarities in data between sampling years and similar patterns observed in the O horizon samples, for which the protocol remained consistent between years.) The three samples were combined by horizon at each site and mixed gently by hand in a bag. From these site-level samples, sub-samples were collected for microbial community analysis and stored in LifeGuard Soil Preservation solution (QIAGEN, Germantown, MD) in a 5 mL tube (Eppendorf, Hamburg, Germany). Tubes were kept as cold as possible while in the field (usually for less than 8 h, but up to 2 days for remote sites) and then stored frozen. The remaining soil samples were air-dried and analyzed for a range of properties (Whitman et al., 2019). Both pH (1:2 soil:DI water for mineral samples; 1:5 soil:DI water for organic samples) and total C (combustion analysis) were measured in both years; soil texture was only measured one year post-fire.

### 2.3 DNA extraction, amplification, and sequencing

DNA extractions were performed for each sample, with two blank extractions for every 24 samples (identical methods but using empty tubes, half of which were sequenced), using a DNeasy PowerLyzer PowerSoil DNA extraction kit (QIAGEN, Germantown, MD) following manufacturer’s instructions. (For the one year post-fire samples, duplicate DNA extractions were performed and sequenced. Duplicates were highly similar, so single extractions were performed for the five years post-fire samples.) Extracted DNA was amplified in triplicate PCR, targeting the 16S rRNA gene v4 region (henceforth, “16S”) with 515f and 806r primers (Walters et al., 2015) with barcodes and Illumina sequencing adapters added as per (Kozich et al., 2013) (all primers in Supplemental Table S1). The PCR amplicon triplicates were pooled, purified and normalized using a SequalPrep Normalization Plate (96) Kit (ThermoFisher Scientific, Waltham, MA). Samples, including blanks, were pooled and library cleanup was performed using a Wizard SV Gel and PCR Clean-Up System A9282 (Promega, Madison, WI). The pooled library was submitted to the UW Madison Biotechnology Center (UW-Madison, WI) for 2×250 paired end (PE) Illumina MiSeq sequencing for the 16S amplicons.

### 2.4 Sequence data processing and taxonomic assignments

We quality-filtered and trimmed, dereplicated, learned errors, determined operational taxonomic units (OTUs), and removed chimeras from 16S reads using dada2 (Callahan et al., 2016) as implemented in R. These sequence processing steps were performed on the UW-Madison Centre for High Throughput Computing cluster (Madison, WI). Taxonomy was assigned to the 16S reads using a QIIME2 (Bolyen et al., 2019) scikit-learn feature classifier (Bokulich et al., 2018) trained on the 515f-806r region of the 99% ID OTUs from the Silva 119 database (Pruesse et al., 2007; Quast et al., 2013; Yilmaz et al., 2013). Our primers target domain *Archaea* as well as domain *Bacteria*, but archaea made up only a small fraction of all sequences (mean across samples <0.1%), so for simplicity, we use the term “bacteria” throughout, rather than “bacteria and archaea”. We predicted 16S rRNA gene copy numbers using the ribosomal RNA operon database (rrnDB) (Stoddard et al., 2015). Because the OTU clustering algorithm can distinguish taxa that differ by a single nucleotide and we used identical sequencing and bioinformatics protocols between the two years, we were able to merge the two datasets for concurrent analysis where necessary. Where we compare five-year data to our one year post-fire dataset, we do all analyses with only the sites that were included in both years of sampling, so minor differences (but no qualitative differences) are to be expected between the one-year data presented here and in the original paper (Whitman et al., 2019).

### 2.5 Co-occurrence networks

To determine which OTUs co-occurred across samples, we used a network analysis approach as in Whitman et al. (2019), following Connor et al. (2017) to avoid false positives and establish conservative network cutoff parameters. After simulating a null model network to choose an appropriate rho value, we determined a consensus network by adding random tie-breaking noise to the matrix 1000 times, selecting only the co-occurrences that occurred in 95% of the 1000 replications. We determined standard network characterization metrics (Guimera and Amaral, 2005; Oleson et al., 2010; Zhou et al., 2010; Deng et al., 2012; Shi et al., 2016), including modularity using random walks, and plotted the network using *igraph* R package (Csardi and Nepusz, 2019). We created networks independently for each year of data. We also note that it is important to limit interpretation of co-occurrence networks, recognizing that co-occurrences do not necessarily represent ecological interactions, such as competition, mutualisms, or predation – rather, they simply indicate whether two organisms tend to co-occur in the same environments.

### 2.6 Statistical analyses

All analyses and plotting were done with R (Team, 2022) in RStudio, using packages *phyloseq* (McMurdie and Holmes, 2013), *dplyr* (Wickham et al., 2021), and *ggplot2* (Wickham, 2016).

To determine whether the same site and soil sample parameters remained significant predictors of community composition five years post-fire, we ran a permutational multivariate ANOVA (PERMANOVA) on Bray-Curtis dissimilarities (Bray and Curtis, 1957) with Hellinger-transformed relative abundances using the *vegan* package in R (Oksanen et al., 2021), reporting R^2^ and p-values for each year’s dataset.

To determine whether burned communities were more similar to unburned communities five years post-fire *vs*. one year post-fire, we compared Bray-Curtis dissimilarities on Hellinger-transformed relative abundances between the two years using a Mann-Whitney U test (because dissimilarities one year post-fire were not normally distributed). We plotted the dissimilarities using non-metric multidimensional scaling (NMDS). To determine whether community composition of severely burned sites changed more between one and five years post-fire, we tested whether Bray-Curtis dissimilarities on Hellinger-transformed relative abundances for paired one-year and five-year samples from the same site were significantly correlated with burn severity index using a Kruksal-Wallis test.

In order to identify which OTUs were significantly enriched (“positive response”) or depleted (“negative response”) in burned plots (*vs*. unburned plots) for each year individually, we used metagenomeSeq (Paulson et al., 2013), after controlling for (including as variables) vegetation community (categorical variable), pH (continuous variable), and %C (continuous variable), resulting in an estimate of the log_2_-fold change in the abundance of each OTU in burned vs. unburned plots, across samples. To determine whether the magnitude of positive or negative responses to fire decreased five years post-fire, we compared the log_2_-fold change for taxa that responded significantly and in the same direction both years using a Mann-Whitney U test, since data were non-normally distributed.

We calculated the abundance-weighted mean predicted copy number for each sample using the approach of Nemergut et al. (2016). We note that this approach is limited in that taxa without representatives in the rRNA gene copy number database will not be represented. We also stress that, while rRNA gene copy number is often correlated with early-successional communities or fast-growing taxa, there are, of course, numerous factors that would also be expected to control fast growth potential, not least of which would be environmental conditions. To determine whether weighted mean predicted rRNA gene copy number was significantly different five years *vs*. one year post-fire, we used a paired Wilcoxon signed rank test, since data were not normally distributed (Shapiro-Wilk test, p<0.05). We also tested whether weighted mean predicted rRNA gene copy number was still significantly correlated with burn severity five years post-fire using a Kruksal-Wallis test.

## 3. Results

### 3.1 Theme 1. Microbial community composition

Five years post-fire, vegetation community, moisture regime, pH, total C, texture, and burned/unburned all remained significant predictors of microbial community composition with similar predictive value (R^2^) (Table 1), although the R^2^ for burned/unburned decreased from 0.07 to 0.04, and vegetation community R^2^ increased from 0.07 to 0.10.

**Table 1.** PERMANOVA results one year (2015) and five years (2019) post-fire. Parameters are listed in order of inclusion in model.

| Parameter | $R^2$ | | p-value | |
| --- | --- | --- | --- | --- |
|  | 2015 | 2019 | 2015 | 2019 |
| Vegetation Community | 0.07 | 0.10 | 0.001 | 0.001 |
| Moisture Regime | 0.02 | 0.02 | 0.002 | 0.030 |
| pH | 0.10 | 0.10 | 0.001 | 0.001 |
| Total C (%) | 0.04 | 0.05 | 0.001 | 0.001 |
| Sand (%) | 0.03 | 0.02 | 0.002 | 0.023 |
| Burned / Unburned | 0.07 | 0.04 | 0.001 | 0.001 |
| Residuals | 0.67 | 0.68 |  |  |

Five years post-fire, the overall structure of the co-occurrence network remained broadly similar (Figures 2a and 2b; Supplemental Table S2). Numerous taxa and co-occurrences were detected in both timepoints: 151 out of 237 total taxa (nodes) in the network one year post-fire were also included in the network five years post-fire (which has 350 total taxa), and 299 out of 1283 edges and their directionality (positive or negative co-occurrences) were maintained. While most pairs of taxa maintained their same co-occurrence patterns (positive or negative in both years), we did see a subset of taxa that shifted from positive co-occurrence to negative co-occurrence (7 pairs, or 21% of sign flips), or vice versa (27 pairs, or 79% of sign flips).

**Figure 2.**
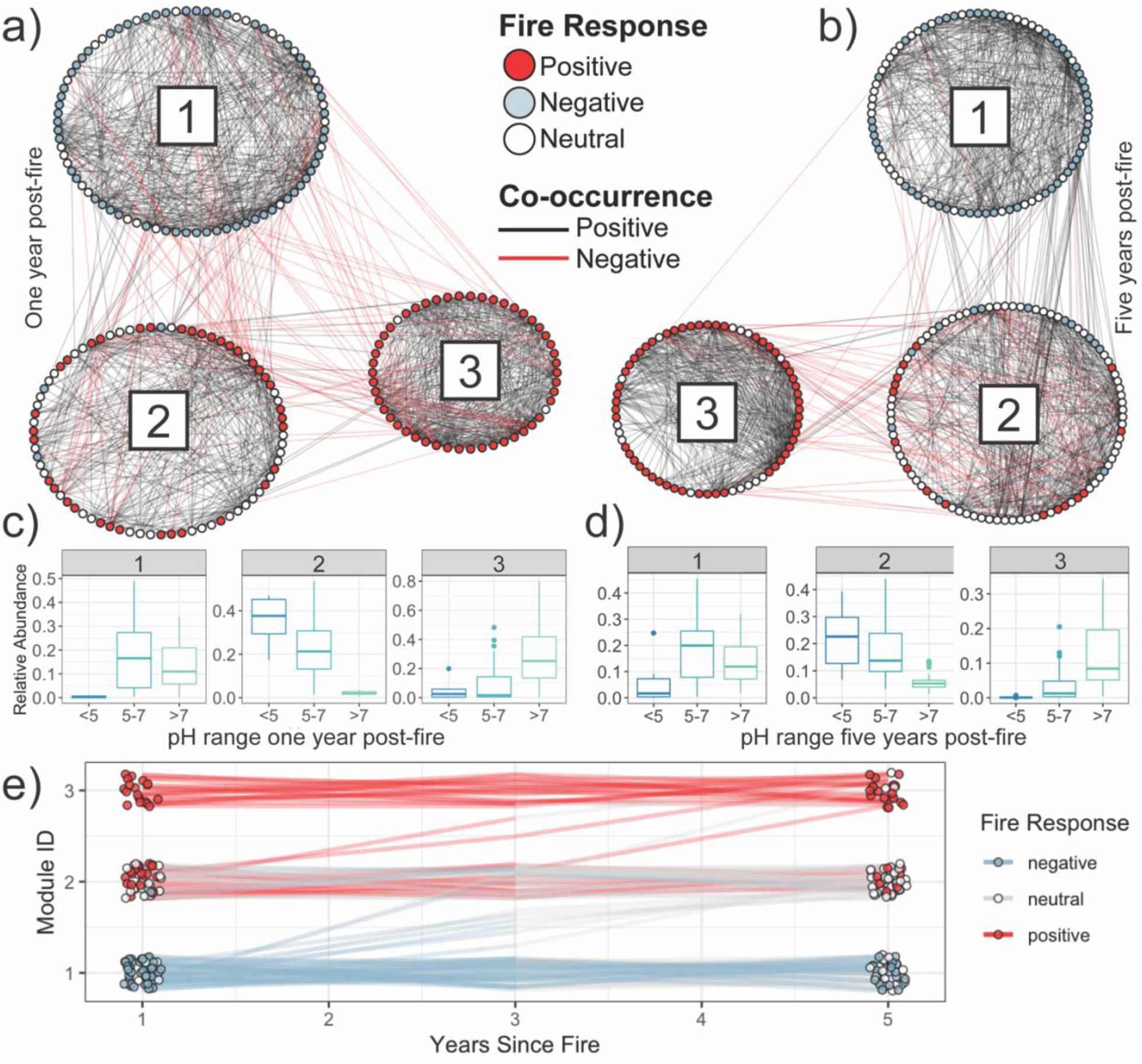
(a and b) The three largest modules of bacterial co-occurrence networks one year (a) and five years (b) post-fire. Each node represents an individual OTU. OTUs joined by edges represent significant co-occurrences (black) or co-exclusions (red). OTUs are clustered into modules of co-occurring taxa, with only the three most abundant modules for each year represented in the figures. OTUs are coloured by response to fire as determined by differential abundance (negative = blue, neutral or not identified = white, positive = red). (c and d) Relative abundance of taxa from a given module across sites within a given pH range, for networks created one year (c) and five years (d) post-fire. (e) Module composition in co-occurrence networks generated independently one year and five years post-fire. Each point represents a single OTU and is coloured by its response to fire within a given year (negative = blue, neutral or not identified = white, positive = red). The same OTU in the two different years is joined by a line, which is also split-coloured by the fire response for the two years. Only OTUs that were present both years in the three most abundant modules are represented.

The same taxa tended to co-occur and cluster into equivalent modules in both years (Figures 2c, 2d, and 2e; Supplemental Figure S1). One year post-fire, the three dominant modules were characterized by taxa mostly negatively associated with fire (Module 1), taxa mostly positively associated with fire and lower pH (Module 2), and taxa almost all positively associated with fire and higher pH (Module 3). Five years post-fire, taxa from the previous fire-responsive, high pH Module 3 tended to remain clustered, with mostly fire-responsive taxa, although the module now contains taxa that are abundant at sites with a broader range of pH values. Taxa from the moderately fire-responsive, low pH Module 2 were associated primarily with new Module 2, and were less likely to be identified as being enriched at burned sites. Many taxa within the negative fire responder Module 1 remained in the new negative fire responder Module 1.

Five years post-fire, bacterial and archaeal communities in burned sites had become more similar to those of unburned sites than they were one year post-fire (Figure 3a; Mann-Whitney U test, p<0.0001). The degree to which they became more similar was not different across severity levels (Figure 3b; Kruksal-Wallis test, p=0.19).

**Figure 3.**
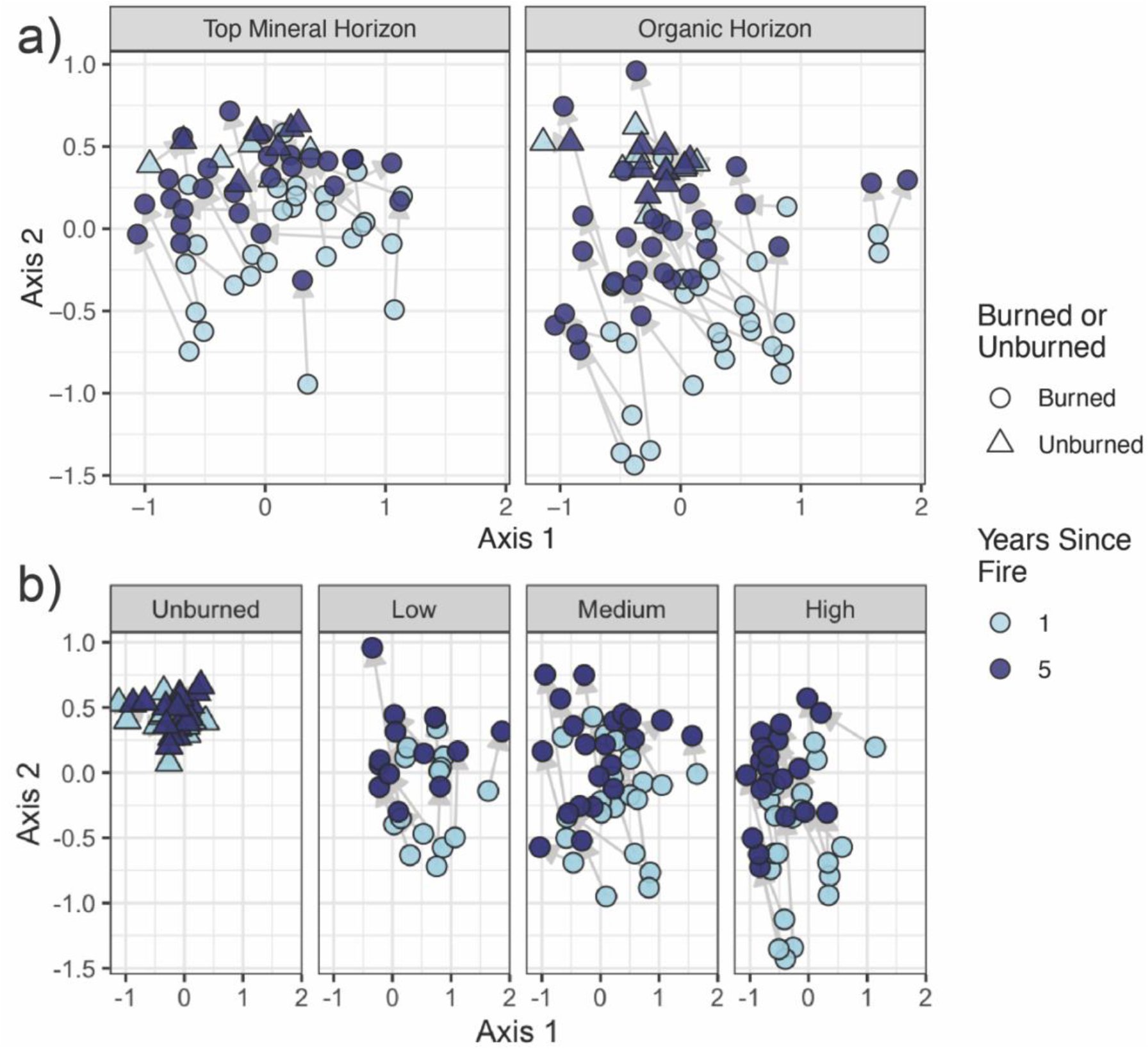
Non-Metric Multidimensional Scaling (NMDS) plots of Bray-Curtis dissimilarities between communities on Hellinger-transformed relative abundances (k=3, stress=0.12). Plots are faceted to illustrate trends, but the panels all draw from the same ordination. Each point represents a single community. Circles represent burned sites, while triangles represent unburned sites, pale blue points were sampled one year post-fire, while dark blue points were sampled five years post-fire, and light grey arrows link the same site between years. (a) Illustrating mineral (M) and organic (O) horizons. (b) Illustrating burn severity categories.

### 3.2 Theme 2. Fire-responsive taxa and traits

Five years post-fire, weighted mean predicted rRNA gene copy number declined significantly (paired t-test, p<0.0001; Figure 4d), and was no longer positively correlated with burn severity index (Kruksal-Wallis test, p=0.23).

**Figure 4.**
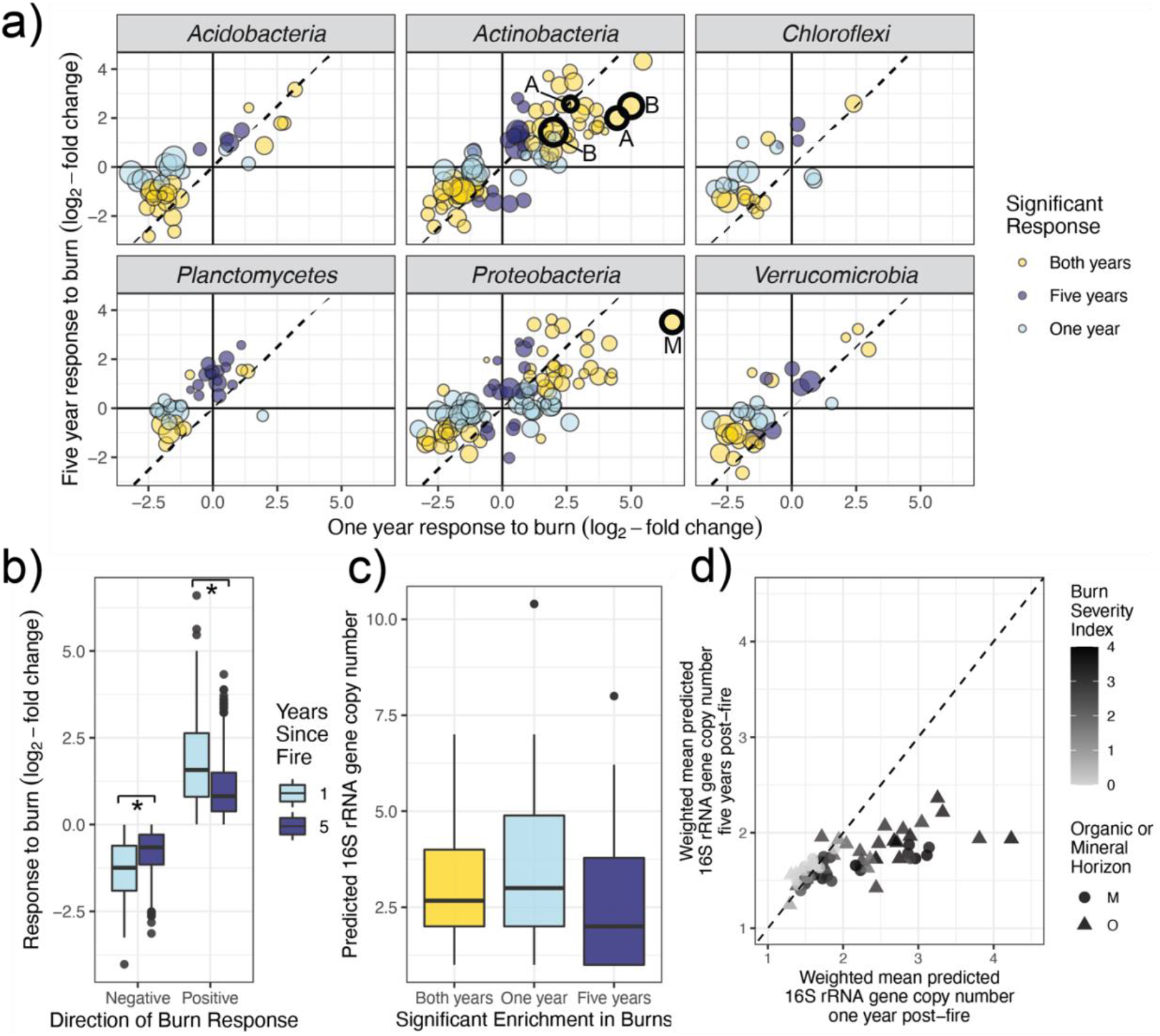
(a) Log_2_-fold change in burned vs. unburned sites one year post fire (x-axis) vs. five years post-fire (y-axis) for significant responders in selected phyla. Dashed line represents 1:1. Each point represents a single OTU. Points are scaled by their mean relative abundance five years post-fire. Points are coloured by whether the response was significant in a given year – yellow indicates significance both years, dark blue represents significant response only five years after fire, and light blue represents significant response only one year after fire. Taxa of interest from the genera *Massilia* (M), *Blastococcus* (B), and *Arthrobacter* (A) have thicker outlines. (b) Response to fire from positive and negative responders one year after fire (light blue) and five years after fire (dark blue). (c) Predicted 16S rRNA gene copy number for OTUs classified as positive fire-responders one, five, or both years after fire. (d) Weighted mean predicted 16S rRNA gene copy number one year post-fire vs. five years post-fire. Points are shaded by burn severity index, with triangles indicating organic horizons (O) and circles indicating mineral horizons (M). Dashed line indicates 1:1 ratio.

Five years post-fire, we found more enriched (339 *vs*. 197 one year post-fire) and a similar number of depleted taxa (245 *vs*. 214 one year post-fire) in burned vs. unburned sites. Responsive taxa generally had similar responses for both years (Figure 4a, yellow points; Supplemental tables S3 and S4), although the responses of individual taxa were also generally muted (less positive or less negative) five years after the fire, compared to one year post-fire (Figure 4b; Mann-Whitney U tests, p=0.009 for positive responders, p<0.0001 for negative responders), and there were numerous individual exceptions, where taxa were enriched or depleted five years post-fire but not one year post-fire (Figure 4a, dark blue points), or were enriched or depleted one year post-fire, but not four years later (Figure 4a, light blue points). The three taxa for which we had specific hypotheses – *Arthrobacter, Massilia*, and *Blastococcus* (heavy black outlines in Figure 4a) – were significantly enriched in burned sites both years. Taxa that were enriched in burned sites one, five, or both years post-fire did not have significantly different predicted rRNA gene copy numbers (Figure 4c; Mann-Whitney U test, p=0.53).

## 4. Discussion

### 4.1 Theme 1. Similar factors control bacterial community composition between one and five years after fire

Consistent with our expectations, we found that the same predictors of bacterial community composition remained significant (Table 1), with similar or even identical R^2^ values for most of them (pH, sand %, C %, and moisture regime, but not burned/unburned, which we discuss in detail later). The exception was the vegetation community, for which R^2^ increased. The increase in the predictive power of vegetation community between one and five years post-fire is consistent with some of the ideas presented in Hart et al. (2005): this system has a relatively long natural fire-return interval, which would indicate that the vegetation-mediated impact of fire on microbial communities may be more driven by specific vegetation composition than nutrient mineralization, particularly over the long term. The importance of the vegetation community in determining the characteristics and regeneration rate of depleted or even completely combusted O horizons would increase over time as this process takes place. As enumerated by Hart et al. (2005), after a wildfire, different plant communities may have different effects on (1) resource inputs via net primary productivity, (2) competition for nutrients, (3) nutrient chemistry, and (4) specific mutualisms (or other ecological interactions), all with implications for the post-fire microbial community.

These findings are also consistent with our second prediction – the general structure of the co-occurrence network remained very similar (Figure 2). Since the factors that structure microbial communities remained constant, then it is reasonable to expect that these same factors are likely still determining which taxa positively or negatively co-occur, resulting in similar co-occurrence networks. We noted with interest the degree to which the networks remained similar. The recreation of this very similar network using all new samples from the same sites, four years after the first sampling, is encouraging, in that it indicates that the methods used to construct the network are robust, and may further strengthen our confidence that its structure is meaningful, rather than statistical noise. That is, the ecological niches of the bacteria observed in this study appear to be broadly consistent for the first several years after fire, in that they still tend to occur in similar environments.

Beyond broad network similarity, we had predicted that the five years post-fire network might have fewer nodes and connections, since we had expected that all sites might begin to converge back to their original unburned composition, thereby decreasing dissimilarity within the dataset and decreasing our ability to detect significant co-occurrences. Although we did see burned sites become more similar to unburned sites five years post-fire, this prediction was not borne out – in fact, the five-year network had more nodes and edges than the one-year network. On the one hand, this could suggest that either the co-occurrences and co-exclusions are sufficiently robust that they remained detectable, even as sites began to return to their pre-burned state. On the other hand, this could be due to the emergence of (*i.e*., ability to detect) new co-occurrences and co-exclusions in the later dataset. The reality is likely a combination of both factors. We do observe that almost a quarter of the co-occurrences (edges) were consistent between both years, supporting the first explanation. However, we also observed the emergence of new co-occurrences. This could occur as new ecological niches are (re-)formed as the soil system recovers post-burn, or as new taxa arrive or increase in abundance at the sites, whether from *in situ* recovery, or due to dispersal from elsewhere (*e.g*., from lower soil horizons, adjacent sites, or airborne particulates (Kobziar et al., 2018)).

While we correctly predicted that burned/unburned would remain a significant explanatory factor for bacterial community composition, counter to our expectations, the R^2^ of burned/unburned for predicting bacterial composition decreased from 0.07 to 0.04 (Table 1) between one and five years post-fire. Even though we did expect to see evidence for resilience (1c), based on chronosequence studies, we had predicted that the four additional years of recovery captured in this study would not meaningfully decrease the predictive power of the burn. Our expectation was that some of the fire-induced changes would take time to emerge, particularly those mediated by vegetation, so burned/unburned would remain an important predictor. However, it seems from our findings that while this effect does seem to be occurring, it may be manifesting as an increased influence of vegetation community (R^2^ of vegetation community as a predictor in the PERMANOVA increased), rather than a continued influence of simply whether or not a site was burned.

At the whole-community level, we generally observed resilience to fire – *i.e*., five years post-fire, communities in burned sites resembled those of unburned sites significantly more than they did one year after the fires (Figure 3a). Furthermore, even severely burned sites recovered as much as low-severity sites did (Figure 3b). That said, the effects of burning remain clearly evident, as would be predicted from chronosequence studies (Sun et al., 2015; Dove and Hart, 2017). Some of these differences are likely to persist for many years because the soil environment, in some sites, remains effectively gone – for example, at sites where the O horizon has largely been combusted. We might expect that the return of the bacterial community to these sites would mirror the redevelopment of the O horizon over time. Other soil properties that were affected by fire may also return to pre-burn conditions relatively slowly. For example, pH can remain altered at burned sites for decades after wildfires (Dymov et al., 2018). Since pH is one of the strongest predictors of microbial community composition in this and other studies (Ramirez et al., 2014; Bahram et al., 2018), it is reasonable to expect that some of the effects of the burns will remain detectable over the same timescales that pH remains significantly altered. While significant differences remain, for reasons such as those just discussed, the broad trend is one of recovery. This may be due to at least two factors. First, some soil properties generally return to their previous state relatively quickly after fires, such as mineral nitrogen pools (Certini, 2005). Second, the return or recolonization of locally-depleted microbial taxa may occur relatively rapidly, whether through dispersal from less-burned patches aboveground or from less-affected soils below the surface, where temperatures from even extreme burns may not reach lethal levels (Pingree and Kobziar, 2019). Overall, our findings suggest that the bacterial community is on a resilient trajectory, following the wide range of burns considered in this study, which includes high severity burns typical of the historical fire regime in this region. However, it is important to note that these fire regimes are characterized by patchiness and spatial variability of fire occurrence and severities (Coop et al., 2020). Depending on the relative importance of post-fire recolonization mechanisms, a shift toward more frequent or more severe wildfires could challenge the resilience observed in this study. For example, if aboveground sources of microbes are critical for recolonization, large or widespread fires could reduce the availability of recolonizing populations, as is observed with the effects of high severity fires on refugia as a source of recolonizing plant seeds (Coop et al., 2019, 2020). If belowground sources are critical for recolonization, fires that reach higher temperatures, resulting in microbial mortality within deeper soil horizons, could also reduce the availability of recolonizing populations, as is observed with the effects of high severity fires on vegetative banks as a source of recolonizing plant materials (Lee, 2004). Thus, the relative importance of *in situ* recovery of surviving taxa and recolonization of burned soils from both above and below for structuring post-fire communities is an area ripe for future investigation. Research on ectomycorrhizal soil fungi after wildfires has indicated that much of post-fire colonization occurs from taxa present in the soil, rather than those dispersed aerially from unburned stands (BAAR et al., 1999; Peay et al., 2009; Glassman et al., 2016). Expanding such work to soil bacterial communities and with an emphasis on differences in wildfire severity would be of interest.

### 4.2 Theme 2. The importance of different fire-responsive traits changes during post-fire recovery

Five years post-fire, predicted fast growth potential was no longer an important factor structuring bacterial communities (Figure 4d). Despite a strong positive correlation with burn severity one year post-fire, at the five-year timepoint, weighted mean predicted rRNA copy number declined significantly overall and was no longer significantly correlated with burn severity. While we had predicted a decline in the strength of this relationship, we had expected that it would remain statistically significant, which was not the case. Our results are consistent with the observations of Nemergut et al. (2016), who observed a significant decline in predicted mean rRNA gene copy number between three months and just over two years post-fire. However, their research was in a temperate ecosystem, while this study took place in the boreal forest. We had thought that perhaps colder conditions overall might slow down the rate of succession as fast growers became less dominant members of the community, but this does not seem to be the case. Reconsidering this idea, it is conceivable that the cool boreal conditions actually decrease the ecological value of fast growth potential, perhaps in favour of other microbial traits, such as stress tolerance. In boreal environments, stress tolerance could be relevant for surviving the heat of the fire, persisting under increased UV exposure (for microbes now at the exposed mineral surface), and tolerating cold conditions. Studying how the timing of post-fire successional trajectories varies for microbial communities across different biomes would be an interesting future focus.

Although community-level predicted mean rRNA gene copy number decreased between one and five years post-fire, we did not see a significant decrease in rRNA copy number among responsive taxa between the two years (Figure 4c). This suggests that even though it is less important at a whole-community level, the enriching effect of an early fast growth strategy may persist for years after the fire for certain taxa. Alternately, it could indicate taxa that possess multiple adaptive post-fire strategies – e.g., some fast-growing taxa might also be particularly well adapted to post-fire environmental conditions.

Our broad hypotheses about fire-responsive taxa were supported (Figures 4a and 4b). Five years post-fire, many taxa remained enriched, some new taxa became enriched, and some taxa were no longer enriched. We also found that 1.46% and 1.75% of all OTUs were enriched one and five years post-fire, respectively, which was consistent with our prediction that a similar number would remain enriched. These findings identify taxa with different timescales of response to fire. Those that became newly enriched five years post-fire might include well-adapted but slow growing taxa and taxa responding to emerging conditions, including taxa that were locally depleted post-fire and needed to recolonize from above or below. Those that were only enriched one year post-fire might indicate taxa that are more classically “ruderal” – quickly growing to take advantage of freed-up niche space and readily available nutrients, but are out-competed as resources become more scarce and competition pressure increases. Taxa enriched both years may be particularly well-adapted to fire and might be more likely to possess multiple fire-adaptive traits. Of the responsive taxa, responses tended to be largest one year post-fire (Figure 4b). However, this was a relatively small decrease – there were still numerous taxa that were markedly enriched (and depleted) compared to unburned sites five years post-fire.

Our fire-responders of interest (*Blastococcus, Arthrobacter*, and *Massilia*) all remained significantly enriched five years post-fire (black circles in Figure 4a). While there were small increases or decreases in the enrichment of each of these taxa, they generally remained enriched to similar degrees. *E.g*., while *Massilia* decreased in its relative enrichment, it remained one of the most fire-enriched bacteria in the study. We had hypothesized that the response from *Blastococcus* might be comparatively short-lived, which we did not observe. This suggests that these *Blastococcus* OTUs could either be remaining enriched due to traits that allowed them to survive or proliferate shortly after the fires, or that they may also have traits that allow them to flourish in the post-fire environment specifically (*e.g*., tolerance to increased pH or ability to degrade PyOM). These hypotheses would need to be tested directly in future studies. Additionally, it would be interesting to compare the timescale of response of these widespread fire-responsive taxa in other different ecosystems – what seems like a persistent effect on *Blastococcus* in the boreal forest could be shorter-lived in warmer systems with more active microbial communities.

We had expected that different traits would change in their relative importance over time since fire. Immediately after fire, survival would be the most important, at least for high-severity burns with significant mortality, while fast growth would emerge next as an important trait. Fast growers might be partially seeded through fire survivors, but also via taxa that recolonize from soils below that experienced sub-lethal temperatures, and from above, via dispersal. Over time, the importance of both of these traits would fade, as the more important traits would be adaptations to the post-fire environment, such as ability to grow at a higher pH or ability to mineralize PyOM. Our data here did not allow us to test the first trait (fire survival). However, they are consistent with a waning influence of the second (fast growth). The third “trait” (adaptation to the post-fire environment) actually encompasses multiple traits.

Taxa that thrive under the chemical conditions that exist post-fire (and may or may not also be fire survivors and fast growers) may dominate the community over longer timescales, to the extent that those chemical conditions also persist over longer timescales. For example, chemical conditions such as increased mineral N (often characterized as a pulse of NO_3_^-^ followed by a pulse of NH_4_^+^) may persist only for a few years, whereas other chemical conditions such as increased pH may persist over intermediate timescales (Certini, 2005; Certini et al., 2021). The ability to metabolize PyOM could remain relevant over long timescales, since PyOM tends to be persistent. However, PyOM degradation might initially be an important trait as a method of detoxification of the environment (removing toxic polyaromatic hydrocarbons), with its relevance as a meaningful source of C emerging within some intermediate timescale – *e.g*., after easily mineralizable fire-liberated C in the environment is depleted, but perhaps before abundant new C has been replenished by regenerating vegetation. Of the numerous traits that may support adaptation to post-fire environments, ability to tolerate pH shifts may be one of the most important ones. This is supported by the data illustrated in Figure 2, which identifies specific taxa in the co-occurrence network Module 3 that remain enriched in burned sites both one and five years post-fire (Figure 2a and 2b), and tend to be more abundant in sites with higher pH (Figure 2c and 2d). Furthermore, pH remains a strong predictor of community composition both one and five years post-fire (Table 1).

Finally, it is essential to note that our ability to conclusively test for the presence and expression of these traits remains limited given the techniques used here (amplicon sequencing amongst a dearth of cultured isolates), and so our findings at this point should serve more as hypothesis generators, to be tested using tools such as shotgun metagenomics, metatranscriptomics, and metaproteomics (illuminating the presence of genes related to potential fire-related traits, their expression, and translation into proteins, respectively), complemented by continued efforts to isolate fire-responsive taxa and measure specific traits in the lab or field. Thoroughly understanding the resilience of bacterial communities and their functions after wildfires will ultimately encompass bacterial community ecology (Hawkes and Keitt, 2015), interactions with plants and fungi, and further investigation of spatial and temporal dynamics, such as recolonization from sub-soils or aerially deposited microbes (Kobziar et al., 2018). These approaches will need to be considered across different ecosystems, fire regimes, and changing climatic conditions.

## Acknowledgements

Thanks to Marc-André Parisien, Mike D. Flannigan, Daniel K. Thompson, Denyse Dawe, and Élyse Mathieu for their contributions in the field and/or to the original study design. This work was supported by the Government of the Northwest Territories, which provided in-kind and financial support for the field campaign that produced these data; Parks Canada Agency and Jean Morin provided in-kind support during fieldwork; the U.S. Department of Energy helped support T.W. and J.W. during data collection [DE-SC0016365; DE-SC0020351] and the U.S. National Science Foundation helped support T.W. during data analysis [2045864].

## CRediT Author Statement

**Thea Whitman:** Conceptualization, Methodology, Software, Formal Analysis, Investigation, Resources, Writing – Original Draft, Visualization, Supervision, Project Administration, Funding Acquisition; **Jamie Woolet:** Investigation, Writing – Review & Editing, Project Administration; **Miranda Sikora:** Formal Analysis, Investigation, Writing – Review & Editing; **Dana Johnson:** Investigation, Writing – Review & Editing; **Ellen Whitman:** Conceptualization, Methodology, Investigation, Resources, Writing – Review & Editing, Visualization, Supervision, Project Administration, Funding Acquisition

## Data Availability

Code for data analysis is available at https://github.com/TheaWhitman/WoodBuffalo1yr5yr. Sequence reads are deposited in the NCBI SRA for 2015 data (PRJNA564811) and 2019 data (PRJNA825513, available 31 December 2022).

## References

Adkins, J., Docherty, K.M., Gutknecht, J.L.M., Miesel, J.R., 2020. How do soil microbial communities respond to fire in the intermediate term? Investigating direct and indirect effects associated with fire occurrence and burn severity. Science of The Total Environment 745, 140957. doi:10.1016/j.scitotenv.2020.140957

Allison, S.D., Martiny, J.B.H., 2008. Colloquium paper: resistance, resilience, and redundancy in microbial communities. Proceedings of the National Academy of Sciences of the United States of America 105 Suppl 1, 11512–11519. doi:10.1073/pnas.0801925105

Anderson, M.K., Lake, F.K., 2013. California Indian Ethnomycology and Associated Forest Management. Journal of Ethnobiology 33, 33–85. doi:10.2993/0278-0771-33.1.33

Andrieux, B., Beguin, J., Bergeron, Y., Grondin, P., Paré, D., 2018. Drivers of postfire soil organic carbon accumulation in the boreal forest. Global Change Biology 24, 4797–4815. doi:10.1111/gcb.14365

Baar, J., Horton, T.R., Kretzer, A.M., Bruns, T.D., 1999. Mycorrhizal colonization of Pinus muricata from resistant propagules after a stand-replacing wildfire. New Phytologist 143, 409–418. doi:10.1046/j.1469-8137.1999.00452.x

Bahram, M., Hildebrand, F., Forslund, S.K., Anderson, J.L., Soudzilovskaia, N.A., Bodegom, P.M., Bengtsson-Palme, J., Anslan, S., Coelho, L.P., Harend, H., Huerta-Cepas, J., Medema, M.H., Maltz, M.R., Mundra, S., Olsson, P.A., Pent, M., Põlme, S., Sunagawa, S., Ryberg, M., Tedersoo, L., Bork, P., 2018. Structure and function of the global topsoil microbiome. Nature 320, 1. doi:10.1038/s41586-018-0386-6

Beckingham, J., Archibald, J., 1996. Field guide to ecosites of Northern Alberta. Natural Resources Canada, Canadian Forest Service, Northern Forestry Centre.

Bender, E.A., Case, T.J., Gilpin, M.E., 1984. Perturbation Experiments in Community Ecology: Theory and Practice. Ecology 65, 1–13.

Bokulich, N.A., Kaehler, B.D., Rideout, J.R., Dillon, M., Bolyen, E., Knight, R., Huttley, G.A., Caporaso, J.G., 2018. Optimizing taxonomic classification of marker-gene amplicon sequences with QIIME 2’s q2-feature-classifier plugin. Microbiome 6, 90. doi:10.1186/s40168-018-0470-z

Bolyen, E., Rideout, J.R., Dillon, M.R., Bokulich, N.A., Abnet, C.C., Al-Ghalith, G.A., Alexander, H., Alm, E.J., Arumugam, M., Asnicar, F., Bai, Y., Bisanz, J.E., Bittinger, K., Brejnrod, A., Brislawn, C.J., Brown, C.T., Callahan, B.J., Caraballo-Rodríguez, A.M., Chase, J., Cope, E.K., Silva, R.D., Diener, C., Dorrestein, P.C., Douglas, G.M., Durall, D.M., Duvallet, C., Edwardson, C.F., Ernst, M., Estaki, M., Fouquier, J., Gauglitz, J.M., Gibbons, S.M., Gibson, D.L., Gonzalez, A., Gorlick, K., Guo, J., Hillmann, B., Holmes, S., Holste, H., Huttenhower, C., Huttley, G.A., Janssen, S., Jarmusch, A.K., Jiang, L., Kaehler, B.D., Kang, K.B., Keefe, C.R., Keim, P., Kelley, S.T., Knights, D., Koester, I., Kosciolek, T., Kreps, J., Langille, M.G.I., Lee, J., Ley, R., Liu, Y.-X., Loftfield, E., Lozupone, C., Maher, M., Marotz, C., Martin, B.D., McDonald, D., McIver, L.J., Melnik, A.V., Metcalf, J.L., Morgan, S.C., Morton, J.T., Naimey, A.T., Navas-Molina, J.A., Nothias, L.F., Orchanian, S.B., Pearson, T., Peoples, S.L., Petras, D., Preuss, M.L., Pruesse, E., Rasmussen, L.B., Rivers, A., Robeson, M.S., Rosenthal, P., Segata, N., Shaffer, M., Shiffer, A., Sinha, R., Song, S.J., Spear, J.R., Swafford, A.D., Thompson, L.R., Torres, P.J., Trinh, P., Tripathi, A., Turnbaugh, P.J., Ul-Hasan, S., Hooft, J.J.J. van der, Vargas, F., Vázquez-Baeza, Y., Vogtmann, E., Hippel, M. von, Walters, W., Wan, Y., Wang, M., Warren, J., Weber, K.C., Williamson, C.H.D., Willis, A.D., Xu, Z.Z., Zaneveld, J.R., Zhang, Y., Zhu, Q., Knight, R., Caporaso, J.G., 2019. Reproducible, interactive, scalable and extensible microbiome data science using QIIME 2. Nature Biotechnology 37, 852–857. doi:10.1038/s41587-019-0209-9

Boudier, M., 1877. De Quelques Espèces Nouvelles De Champignons. Bulletin de la Société Botanique de France 24, 307–312. doi:10.1080/00378941.1877.10830001

Boulanger, Y., Gautier, S., Burton, P.J., Vaillancourt, M.A., 2012. An alternative fire regime zonation for Canada. International Journal of Wildland Fire 21, 1052–1064.

Bowman, D.M.J.S., Balch, J.K., Artaxo, P., Bond, W.J., Carlson, J.M., Cochrane, M.A., D’Antonio, C.M., DeFries, R.S., Doyle, J.C., Harrison, S.P., Johnston, F.H., Keeley, J.E., Krawchuk, M.A., Kull, C.A., Marston, J.B., Moritz, M.A., Prentice, I.C., Roos, C.I., Scott, A.C., Swetnam, T.W., Werf, G.R. van der, Pyne, S.J., 2009. Fire in the Earth System. Science 324, 481–484. doi:10.1126/science.1163886

Bray, J.R., Curtis, J.T., 1957. An Ordination of the Upland Forest Communities of Southern Wisconsin. Ecological Monographs 27, 325–349. doi:10.2307/1942268

Brown, S.P., Veach, A.M., Horton, J.L., Ford, E., Jumpponen, A., Baird, R., 2019. Context dependent fungal and bacterial soil community shifts in response to recent wildfires in the Southern Appalachian Mountains. Forest Ecology and Management 451, 117520. doi:10.1016/j.foreco.2019.117520

Callahan, B.J., McMurdie, P.J., Rosen, M.J., Han, A.W., Johnson, A.J.A., Holmes, S.P., 2016. DADA2: High-resolution sample inference from Illumina amplicon data. Nature Methods 13, 581–583. doi:10.1038/nmeth.3869

Certini, G., 2005. Effects of fire on properties of forest soils: a review. Oecologia 143, 1–10. doi:10.1007/s00442-004-1788-8

Certini, G., Moya, D., Lucas-Borja, M.E., Mastrolonardo, G., 2021. The impact of fire on soil-dwelling biota: A review. Forest Ecology and Management 488, 118989. doi:10.1016/j.foreco.2021.118989

Connor, N., Barberán, A., Clauset, A., 2017. Using null models to infer microbial co-occurrence networks. PLoS ONE 12, e0176751. doi:10.1371/journal.pone.0176751

Coop, J.D., DeLory, T.J., Downing, W.M., Haire, S.L., Krawchuk, M.A., Miller, C., Parisien, M., Walker, R.B., 2019. Contributions of fire refugia to resilient ponderosa pine and dry mixed-conifer forest landscapes. Ecosphere 10. doi:10.1002/ecs2.2809

Coop, J.D., Parks, S.A., Rumann, C.S.S., Crausbay, S.D., Higuera, P.E., Hurteau, M.D., Tepley, A., Whitman, E., Assal, T., Collins, B.M., Davis, K.T., Dobrowski, S., Falk, D.A., Fornwalt, P.J., Fulé, P.Z., Harvey, B.J., Kane, V.R., Littlefield, C.E., Margolis, E.Q., North, M., Parisien, M.-A., Prichard, S., Rodman, K.C., 2020. Wildfire-Driven Forest Conversion in Western North American Landscapes. BioScience 46, 326–15. doi:10.1093/biosci/biaa061

Cooper, C.F., 1961. The Ecology of Fire 204, 150–163.

Csardi, G., Nepusz, T., 2019. The igraph software package for complex network research. InterJournal Complex Systems, 1695.

Czimczik, C.I., Masiello, C.A., 2007. Controls on black carbon storage in soils. Global Biogeochemical Cycles 21, GB3005. doi:10.1029/2006gb002798

Delgado-Baquerizo, M., Oliverio, A.M., Brewer, T.E., Benavent-González, A., Eldridge, D.J., Bardgett, R.D., Maestre, F.T., Singh, B.K., Fierer, N., 2018. A global atlas of the dominant bacteria found in soil. Science 359, 320–325. doi:10.1126/science.aap9516

Deng, Y., Jiang, Y.-H., Yang, Y., He, Z., Luo, F., Zhou, J., 2012. Molecular ecological network analyses. Bmc Bioinformatics 13. doi:10.1186/1471-2105-13-113

Dooley, S.R., Treseder, K.K., 2011. The effect of fire on microbial biomass: a meta-analysis of field studies. Biogeochemistry 109, 49–61. doi:10.1007/s10533-011-9633-8

Dove, N.C., Hart, S.C., 2017. Fire Reduces Fungal Species Richness and In Situ Mycorrhizal Colonization: A Meta-Analysis. Fire Ecology 13, 37–65. doi:10.4996/fireecology.130237746

Dymov, A., Abakumov, E., Bezkorovaynaya, I., Prokushkin, A., Kuzyakov, Y., Milanovsky, E., 2018. Impact of forest fire on soil properties (review). Theoretical and Applied Ecology 827–836.

Eswg, E.S.W.G., 1995. A National Ecological Framework for Canada. Agriculture and Agri-Food Canada, Ottawa, Ontario/Hull, Quebec, Canada.

Fernández-González, A.J., Martínez-Hidalgo, P., Cobo-Díaz, J.F., Villadas, P.J., Martínez-Molina, E., Toro, N., Tringe, S.G., Fernández-López, M., 2017. The rhizosphere microbiome of burned holm-oak: potential role of the genus Arthrobacter in the recovery of burned soils. Scientific Reports 7, 6008. doi:10.1038/s41598-017-06112-3

Glassman, S.I., Levine, C.R., DiRocco, A.M., Battles, J.J., Bruns, T.D., 2016. Ectomycorrhizal fungal spore bank recovery after a severe forest fire: some like it hot. The ISME Journal 10, 1228–1239.

Guimera, R., Amaral, L., 2005. Functional cartography of complex metabolic networks. Nature 433, 895–900. doi:10.1038/nature03288

Hart, S.C., DeLuca, T.H., Newman, G.S., MacKenzie, M.D., Boyle, S.I., 2005. Post-fire vegetative dynamics as drivers of microbial community structure and function in forest soils. Forest Ecology and Management 220, 166–184. doi:10.1016/j.foreco.2005.08.012

Hawkes, C.V., Keitt, T.H., 2015. Resilience vs. historical contingency in microbial responses to environmental change. Ecology Letters 18, 612–625. doi:10.1111/ele.12451

Holden, S.R., Rogers, B.M., Treseder, K.K., Randerson, J.T., 2016. Fire severity influences the response of soil microbes to a boreal forest fire. Environmental Research Letters 11, 10. doi:10.1088/1748-9326/11/3/035004

Holden, S.R., Treseder, K.K., 2013. A meta-analysis of soil microbial biomass responses to forest disturbances. Terrestrial Microbiology 4, 1–17. doi:10.3389/fmicb.2013.00163

Johnstone, J.F., Allen, C.D., Franklin, J.F., Frelich, L.E., Harvey, B.J., Higuera, P.E., Mack, M.C., Meentemeyer, R.K., Metz, M.R., Perry, G.L., Schoennagel, T., Turner, M.G., 2016. Changing disturbance regimes, ecological memory, and forest resilience. Frontiers in Ecology and the Environment 14, 369–378. doi:10.1002/fee.1311

Keeley, J.E., 2009. Fire intensity, fire severity and burn severity: a brief review and suggested usage. International Journal of Wildland Fire 18, 116. doi:10.1071/wf07049

Klappenbach, J.A., Dunbar, J.M., Schmidt, T.M., 2000. rRNA Operon Copy Number Reflects Ecological Strategies of Bacteria. Applied and Environmental Microbiology 66, 1328–1333. doi:10.1128/aem.66.4.1328-1333.2000

Knelman, J.E., Graham, E.B., Trahan, N.A., Schmidt, S.K., Nemergut, D.R., 2015. Fire severity shapes plant colonization effects on bacterial community structure, microbial biomass, and soil enzyme activity in secondary succession of a burned forest. Soil Biology and Biochemistry 90, 161–168. doi:10.1016/j.soilbio.2015.08.004

Kobziar, L.N., Pingree, M.R.A., Larson, H., Dreaden, T.J., Green, S., Smith, J.A., 2018. Pyroaerobiology: the aerosolization and transport of viable microbial life by wildland fire. Ecosphere 9, e02507–12. doi:10.1002/ecs2.2507

Köster, K., Berninger, F., Lindén, A., Köster, E., Pumpanen, J., 2014. Recovery in fungal biomass is related to decrease in soil organic matter turnover time in a boreal fire chronosequence. Geoderma 235–236, 74–82. doi:10.1016/j.geoderma.2014.07.001

Kozich, J.J., Westcott, S.L., Baxter, N.T., Highlander, S.K., Schloss, P.D., 2013. Development of a Dual-Index Sequencing Strategy and Curation Pipeline for Analyzing Amplicon Sequence Data on the MiSeq Illumina Sequencing Platform. Applied and Environmental Microbiology 79, 5112–5120. doi:10.1128/aem.01043-13

Lecomte, N., Simard, M., Fenton, N., Bergeron, Y., 2006. Fire Severity and Long-term Ecosystem Biomass Dynamics in Coniferous Boreal Forests of Eastern Canada. Ecosystems 9, 1215–1230. doi:10.1007/s10021-004-0168-x

Lee, P., 2004. The impact of burn intensity from wildfires on seed and vegetative banks, and emergent understory in aspen-dominated boreal forests. Canadian Journal of Botany 82, 1468–1480. doi:10.1139/b04-108

Lierop, P. van, Lindquist, E., Sathyapala, S., Franceschini, G., 2015. Global forest area disturbance from fire, insect pests, diseases and severe weather events. Forest Ecology and Management 352, 78–88. doi:10.1016/j.foreco.2015.06.010

Liu, J., Liu, S., Sun, K., Sheng, Y., Gu, Y., Gao, Y., 2014. Colonization on Root Surface by a Phenanthrene-Degrading Endophytic Bacterium and Its Application for Reducing Plant Phenanthrene Contamination. PLoS ONE 9, e108249. doi:10.1371/journal.pone.0108249

Loboda, T.V., French, N.H.F., Hight-Harf, C., Jenkins, L., Miller, M.E., 2013. Mapping fire extent and burn severity in Alaskan tussock tundra: An analysis of the spectral response of tundra vegetation to wildland fire. Remote Sensing of Environment 134, 194–209. doi:10.1016/j.rse.2013.03.003

Lucas-Borja, M.E., Miralles, I., Ortega, R., Plaza-Álvarez, P.A., Gonzalez-Romero, J., Sagra, J., Soriano-Rodríguez, M., Certini, G., Moya, D., Heras, J., 2019. Immediate fire-induced changes in soil microbial community composition in an outdoor experimental controlled system. Science of The Total Environment 696, 134033. doi:10.1016/j.scitotenv.2019.134033

Mack, M.C., Walker, X.J., Johnstone, J.F., Alexander, H.D., Melvin, A.M., Jean, M., Miller, S.N., 2021. Carbon loss from boreal forest wildfires offset by increased dominance of deciduous trees. Science 372, 280–283. doi:10.1126/science.abf3903

McMurdie, P.J., Holmes, S., 2013. phyloseq: An R Package for Reproducible Interactive Analysis and Graphics of Microbiome Census Data. PLoS ONE 8, e61217. doi:10.1371/journal.pone.0061217

Miera, L.E.S. de, Pinto, R., Gutierrez-Gonzalez, J.J., Calvo, L., Ansola, G., 2020. Wildfire effects on diversity and composition in soil bacterial communities. The Science of the Total Environment 726, 138636. doi:10.1016/j.scitotenv.2020.138636

Mongodin, E.F., Shapir, N., Daugherty, S.C., DeBoy, R.T., Emerson, J.B., Shvartzbeyn, A., Radune, D., Vamathevan, J., Riggs, F., Grinberg, V., Khouri, H., Wackett, L.P., Nelson, K.E., Sadowsky, M.J., 2006. Secrets of Soil Survival Revealed by the Genome Sequence of Arthrobacter aurescens TC1. PLOS Genetics 2, e214. doi:10.1371/journal.pgen.0020214

Nave, L.E., Vance, E.D., Swanston, C.W., Curtis, P.S., 2011. Fire effects on temperate forest soil C and N storage. Ecological Applications 21, 1189–1201. doi:10.1890/10-0660.1

Nemergut, D.R., Knelman, J.E., Ferrenberg, S., Bilinski, T., Melbourne, B., Jiang, L., Violle, C., Darcy, J.L., Prest, T., Schmidt, S.K., Townsend, A.R., 2016. Decreases in average bacterial community rRNA operon copy number during succession. The ISME Journal 10, 1147–1156. doi:10.1038/ismej.2015.191

Oksanen, J., Blanchet, F.G., Kindt, R., Legendre, P., Minchin, P.R., OHara, R.B., Simpson, G.L., Solymos, P., Stevens, M.H.H., Wagner, H., 2021. vegan: Community Ecology Package.

Oleson, K.W., Lawrence, D.M., Bonan, G.B., Flanner, M.G., Kluzek, E., Lawrence, P.J., Levis, S., Swenson, S.C., Thornton, P.E., Dai, A., Decker, M., Dickinson, R., Feddema, J., Heald, C.L., Hoffman, F., Lamarque, J.-F., Mahowald, N., Niu, G.-Y., Qian, T., Randerson, J., Running, S., Sakaguchi, K., Slater, A., Stockli, R., Wang, A., Yang, Z.-L., Zeng, Xiaodong, Zeng, Xubin, 2010. Technical Description of version 4.0 of the Community Land Model (CLM) [WWW Document]. URL http://www.cesm.ucar.edu/models/ccsm4.0/clm/CLM4_Tech_Note.pdf

Paulson, J.N., Stine, O.C., Bravo, H.C., Pop, M., 2013. Differential abundance analysis for microbial marker-gene surveys. Nature Methods 10, 1200–1202. doi:10.1038/nmeth.2658

Peay, K.G., Garbelotto, M., Bruns, T.D., 2009. Spore heat resistance plays an important role in disturbance-mediated assemblage shift of ectomycorrhizal fungi colonizing Pinus muricata seedlings. Journal of Ecology 97, 537–547. doi:10.1111/j.1365-2745.2009.01489.x

Pérez-Valera, E., Verdú, M., Navarro-Cano, J.A., Goberna, M., 2020. Soil microbiome drives the recovery of ecosystem functions after fire. Soil Biology and Biochemistry 149, 107948. doi:10.1016/j.soilbio.2020.107948

Pérez-Valera, E., Verdú, M., Navarro-Cano, J.A., Goberna, M., 2018. Resilience to fire of phylogenetic diversity across biological domains. Molecular Ecology 27, 2896–2908. doi:10.1111/mec.14729

Pingree, M.R.A., Kobziar, L.N., 2019. The myth of the biological threshold: A review of biological responses to soil heating associated with wildland fire. Forest Ecology and Management 432, 1022–1029. doi:10.1016/j.foreco.2018.10.032

Pressler, Y., Moore, J.C., Cotrufo, M.F., 2018. Belowground community responses to fire: meta-analysis reveals contrasting responses of soil microorganisms and mesofauna. Oikos. doi:10.1111/oik.05738

Pruesse, E., Quast, C., Knittel, K., Fuchs, B.M., Ludwig, W., Peplies, J., Glockner, F.O., 2007. SILVA: a comprehensive online resource for quality checked and aligned ribosomal RNA sequence data compatible with ARB. - PubMed - NCBI. Nucleic Acids Research 35, 7188–7196.

Quast, C., Pruesse, E., Yilmaz, P., Gerken, J., Schweer, T., Yarza, P., Peplies, J., Glöckner, F.O., 2013. The SILVA ribosomal RNA gene database project: improved data processing and web-based tools. Nucleic Acids Research 41, D590–D596. doi:10.1093/nar/gks1219

Ramirez, K.S., Leff, J.W., Barberan, A., Bates, S.T., Betley, J., Crowther, T.W., Kelly, E.F., Oldfield, E.E., Shaw, E.A., Steenbock, C., Bradford, M.A., Wall, D.H., Fierer, N., 2014. Biogeographic patterns in below-ground diversity in New York City’s Central Park are similar to those observed globally. Proceedings. Biological Sciences / The Royal Society 281, 20141988–20141988. doi:10.1098/rspb.2014.1988

Roller, B.R.K., Stoddard, S.F., Schmidt, T.M., 2016. Exploiting rRNA Operon Copy Number to Investigate Bacterial Reproductive Strategies. Nature Microbiology 1, 16160–16160. doi:10.1038/nmicrobiol.2016.160

Seaver, F.J., 1909. Studies in Pyrophilous Fungi: I. The Occurrence and Cultivation of Pyronema. Mycologia 1, 131–139. doi:10.2307/3753124

Shade, A., Peter, H., Allison, S.D., Baho, D.L., Berga, M., Bürgmann, H., Huber, D.H., Langenheder, S., Lennon, J.T., Martiny, J.B.H., Matulich, K.L., Schmidt, T.M., Handelsman, J., 2012. Fundamentals of Microbial Community Resistance and Resilience. Terrestrial Microbiology 3, 417. doi:10.3389/fmicb.2012.00417

Shi, S., Nuccio, E.E., Shi, Z.J., He, Z., Zhou, J., Firestone, M.K., 2016. The interconnected rhizosphere: High network complexity dominates rhizosphere assemblages. Ecology Letters 19, 926–936. doi:10.1111/ele.12630

Smithwick, E.A.H., Turner, M.G., Metzger, K.L., Balser, T.C., 2005. Variation in NH4+ mineralization and microbial communities with stand age in lodgepole pine (Pinus contorta) forests, Yellowstone National Park (USA). Soil Biology and Biochemistry 37, 1546–1559. doi:10.1016/j.soilbio.2005.01.016

Stoddard, S.F., Smith, B.J., Hein, R., Roller, B.R.K., Schmidt, T.M., 2015. rrnDB: improved tools for interpreting rRNA gene abundance in bacteria and archaea and a new foundation for future development. Nucleic Acids Research 43, D593–D598. doi:10.1093/nar/gku1201

Sun, H., Santalahti, M., Pumpanen, J., Köster, K., Berninger, F., Raffaello, T., Asiegbu, F.O., Heinonsalo, J., 2016. Bacterial community structure and function shift across a northern boreal forest fire chronosequence. Scientific Reports 6, 34. doi:10.1038/srep32411

Sun, H., Santalahti, M., Pumpanen, J., Köster, K., Berninger, F., Raffaello, T., Jumpponen, A., Asiegbu, F.O., Heinonsalo, J., 2015. Fungal Community Shifts in Structure and Function across a Boreal Forest Fire Chronosequence. Applied and Environmental Microbiology 81, 7869–7880. doi:10.1128/aem.02063-15

Team, R.C., 2022. R: A language and environment for statistical computing.

Walters, W., Hyde, E.R., Berg-Lyons, D., Ackermann, G., Humphrey, G., Parada, A., Gilbert, J.A., Jansson, J.K., Caporaso, J.G., Fuhrman, J.A., Apprill, A., Knight, R., 2015. Improved Bacterial 16S rRNA Gene (V4 and V4-5) and Fungal Internal Transcribed Spacer Marker Gene Primers for Microbial Community Surveys. MSystems 1, e00009–15. doi:10.1128/msystems.00009-15

Wan, S., Hui, D., Luo, Y., 2001. Fire effects on nitrogen pools and dynamics in terrestrial ecosystems: A meta-analysis. Ecological Applications 11, 1349–1365. doi:10.1890/1051-0761(2001)011[1349:feonpa]2.0.co;2

Wang, T., Hamann, A., Spittlehouse, D.L., Murdock, T.Q., Wang, T., Hamann, A., Spittlehouse, D.L., Murdock, T.Q., 2012. ClimateWNA—High-Resolution Spatial Climate Data for Western North America. Dx.Doi.Org 51, 16–29. doi:10.1175/jamc-d-11-043.1

Weber, C.F., Lockhart, J.S., Charaska, E., Aho, K., Lohse, K.A., 2014. Bacterial composition of soils in ponderosa pine and mixed conifer forests exposed to different wildfire burn severity. Soil Biology and Biochemistry 69, 242–250. doi:10.1016/j.soilbio.2013.11.010

Westerberg, K., Elvang, A.M., Stackebrandt, E., Jansson, J.K., 2000. Arthrobacter chlorophenolicus sp nov., a new species capable of degrading high concentrations of 4-chlorophenol. International Journal of Systematic and Evolutionary Microbiology 50, 2083–2092. doi:10.1099/00207713-50-6-2083

Whitman, E., Parisien, M.-A., Thompson, D., Flannigan, M., 2018a. Topoedaphic and Forest Controls on Post-Fire Vegetation Assemblies Are Modified by Fire History and Burn Severity in the Northwestern Canadian Boreal Forest. Forests 9, 151. doi:10.3390/f9030151

Whitman, E., Parisien, M.-A., Thompson, D.K., Hall, R.J., Skakun, R.S., Flannigan, M.D., 2018b. Variability and drivers of burn severity in the northwestern Canadian boreal forest. Ecosphere 9. doi:10.1002/ecs2.2128

Whitman, T., Whitman, E., Woolet, J., Flannigan, M.D., Thompson, D.K., Parisien, M.-A., 2019. Soil Bacterial and Fungal Response to Wildfires in the Canadian Boreal Forest Across a Burn Severity Gradient. Soil Biology and Biochemistry 138, 107571.

Wickham, H., 2016. ggplot2: Elegant Graphics for Data Analysis, Springer-Verlag New York. Springer-Verlag New York.

Wickham, H., François, R., Henry, L., Müller, K., 2021. dplyr: A Grammar of Data Manipulation.

Yilmaz, P., Parfrey, L.W., Yarza, P., Gerken, J., Pruesse, E., Quast, C., Schweer, T., Peplies, J., Ludwig, W., Glöckner, F.O., 2013. The SILVA and “All-species Living Tree Project (LTP)” taxonomic frameworks. Nucleic Acids Research 42, D643–D648. doi:10.1093/nar/gkt1209

Zhou, J., Deng, Y., Luo, F., He, Z., Tu, Q., Zhi, X., 2010. Functional Molecular Ecological Networks. MBio 1. doi:10.1128/mbio.00169-10

Zhou, X., Sun, H., Sietiö, O.-M., Pumpanen, J., Heinonsalo, J., Köster, K., Berninger, F., 2020. Wildfire effects on soil bacterial community and its potential functions in a permafrost region of Canada. Applied Soil Ecology 156, 103713. doi:10.1016/j.apsoil.2020.103713

## Additional References

Dawe, D., Parisien, M.-A., Van Dongen, A., Whitman, E. Under review at Plant Ecology. Initial Succession After Wildfire in Dry Boreal Forests of Northwestern North America. Preprint at https://www.researchsquare.com/article/rs-1112059/v1.

Woolet, J., Whitman, E., Parisien, M.-A., Thompson, D.K., Flannigan, M., Whitman, T. Under review at FEMS Microbiology Ecology. Short-interval reburns in the boreal forest alter soil bacterial communities compared to long-interval reburns, accompanied by pH shifts and poor conifer seedling establishment. Preprint on BiorXiv at (10.1101/2021.03.31.437944).

