## Supplemental Information for "Resilience in soil bacterial communities of the boreal forest from one to five years after wildfire across a severity gradient"

**Table of Contents**

Supplemental Figure S1. 2015 and 2019 co-occurrence networks 2
Supplemental Table S1. Primers used in this study 3
Supplemental Table S2. Co-occurrence network properties for full networks 4
Supplemental Table S3. Burn-enriched taxa in 2015, one year post-fire 5
Supplemental Table S4. Burn-enriched taxa in 2019, five years post-fire 5

References 6

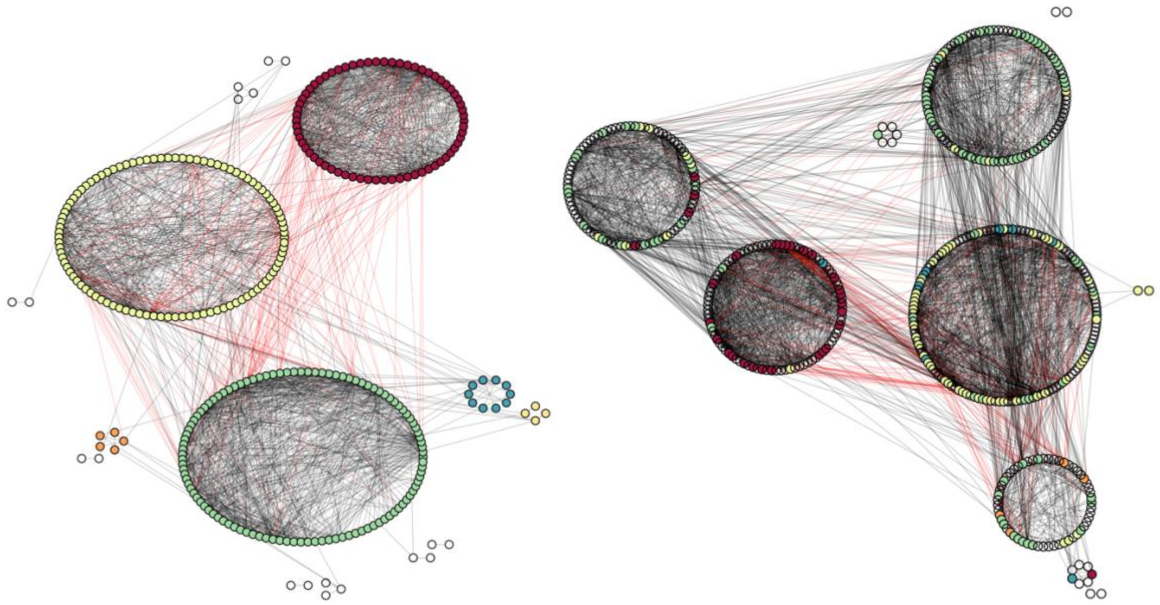

Supplemental Figure S1. Full 2015 (left) and 2019 (right) bacterial co-occurrence networks. Each node represents an individual OTU. OTUs joined by edges represent significant co-occurrences (black) or co-exclusions (red). OTUs are clustered into modules of co-occurring taxa. OTUs are coloured by their module ID in the 2015 network, illustrating moderate correspondence between modules between years (Figure 2 in main manuscript).

Supplemental Table S1. Primers used in this study (Kozich et al., 2013; Walters et al., 2015)

| **Primer** | **Sequence [**Illumina adaptor *Barcode* **Pad and linker** Primer] |
|---|---|
| *16S Illumina* | |
| 515f | AATGATACGGCGACCACCGAGATCTACAC*XXXXXXXX***TATGGTAATTGT**GTGYCAGCMGCCGCGGTAA |
| 806r | CAAGCAGAAGACGGCATACGAGAT*XXXXXXXX***AGTCAGCCAGCC**GGACTACNVGGGTWTCTAAT |
| Read 1 seq | TATGGTAATTGTGTGYCAGCMGCCGCGGTAA |
| Read 2 seq | AGTCAGCCAGCCGGACTACNVGGGTWTCTAAT |
| Barcode seq | ATTAGAWACCCBNGTAGTCCGGCTGGCTGACT |

| Supplemental Table S2. Co-occurrence network properties for full networks | | |
|---|---|---|
| | *One year post-fire* | *Five years post-fire* |
| Rho cutoff value used | 0.43 | 0.43 |
| Total nodes (taxa) | 237 | 350 |
| Total edges (co-occurrences) | 1283 | 1858 |
| Average path length | 4.2 | 3.2 |
| Diameter (longest [shortest path between two nodes]) | 7 | 7 |
| Average degree (mean number of edges at nodes) | 10.8 | 10.6 |
| Edge density | 0.05 | 0.03 |
| Global clustering coefficient (all triangles) | 0.46 | 0.38 |
| Average clustering coefficient (each node) | 0.58 | 0.55 |
| $R^2$ of power-law | 0.68 | 0.72 |
| Exponent (alpha) of power law | 0.89 | 1.02 |
| Modularity | 0.56 | 0.59 |

Supplemental Table S3. Burn-enriched taxa in 2015, one year post-fire
*Enclosed as separate .csv*

Supplemental Table S4. Burn-enriched taxa in 2019, five years post-fire
*Enclosed as separate .csv*

**References**


Kozich, J.J., Westcott, S.L., Baxter, N.T., Highlander, S.K., Schloss, P.D., 2013. Development of a Dual-Index Sequencing Strategy and Curation Pipeline for Analyzing Amplicon Sequence Data on the MiSeq Illumina Sequencing Platform. Applied and Environmental Microbiology 79, 5112–5120. doi:10.1128/aem.01043-13

Taylor, D.L., Walters, W.A., Lennon, N.J., Bochicchio, J., Krohn, A., Caporaso, J.G., Pennanen, T., 2016. Accurate Estimation of Fungal Diversity and Abundance Through Improved Lineage-Specific Primers Optimized for Illumina Amplicon Sequencing. Applied and Environmental Microbiology 82, AEM.02576-16-7226. doi:10.1128/aem.02576-16

Walters, W., Hyde, E.R., Berg-Lyons, D., Ackermann, G., Humphrey, G., Parada, A., Gilbert, J.A., Jansson, J.K., Caporaso, J.G., Fuhrman, J.A., Apprill, A., Knight, R., 2015. Improved Bacterial 16S rRNA Gene (V4 and V4-5) and Fungal Internal Transcribed Spacer Marker Gene Primers for Microbial Community Surveys. MSystems 1, e00009-15. doi:10.1128/msystems.00009-15
